# Cereal Weevil’s Antimicrobial Peptides: At the Crosstalk between Development, Endosymbiosis and Immune Response

**DOI:** 10.1101/2023.09.27.559682

**Authors:** N. Galambos, C. Vincent-Monegat, A. Vallier, N. Parisot, A. Heddi, A. Zaidman-Rémy

## Abstract

Interactions between animals and microbes are ubiquitous in nature and strongly impact animal physiology. These interactions are shaped by the host immune system, which responds to infections and contributes to tailor the associations with beneficial microorganisms. In many insects, beneficial symbiotic associations not only include gut commensals, but also intracellular bacteria, or endosymbionts. Endosymbionts are housed within specialised host cells, the bacteriocytes, and are transmitted vertically across host generations. Host-endosymbiont co-evolution shapes the endosymbiont genome and host immune system, which not only fights against microbial intruders, but also ensures the preservation of endosymbionts and the control of their load and location. The cereal weevil *Sitophilus* spp. is a remarkable model to study the evolutionary adaptation of the immune system to endosymbiosis since its binary association with a unique, relatively recently acquired nutritional endosymbiont, *Sodalis pierantonius*. This Gram-negative bacterium has not experienced the genome size shrinkage observed in long-term endosymbioses and has retained immunogenicity. We focus here on the 16 antimicrobial peptides (AMPs) identified in the *Sitophilus oryzae* genome and their expression patterns in different tissues, along host development or upon immune challenges, to address their potential functions in the defensive response and endosymbiosis homeostasis along the insect life cycle.

## Introduction

Antimicrobial peptides (AMPs) are ancient, broad-spectrum molecules, frequently associated with pathogen clearance ^(1)^. AMPs are produced by a wild variety of organisms ranging from bacteria to humans ^(1,2)^. AMPs can act additively, synergistically or with high specificity against viral, bacterial, fungal and protozoan pathogens ^(1,3–7)^. These small cationic peptides were shown to target microbial phospholipid membranes, leading to depolarisation, changes in their permeability, and subsequent microbial cell death. Some AMPs can also inhibit viral proliferation by disruption of the viral protein synthesis and the viral gene expression ^(1,7,8)^. Although prokaryotic AMPs, called Bacteriocins, have also been described ^(2)^, we will focus here on eukaryotic AMPs.

Research on insect immunity has been at the forefront of antimicrobial peptide studies since the first description of an animal AMP in the silk moth *Hyalophora cecropia* ^(9)^. Cecropin was reported as a novel bactericidal agent able to permeabilise the membrane of a limited number of Gram-positive bacteria, without affecting eukaryotic cells ^(9)^. Since the discovery of this first AMP, a large number of AMPs have been identified in diverse insect families as an important part of the immune response ^(7)^. Interestingly, recent availability of functional analyses is currently uncovering the level of specificity and synergy of these immune effectors ^(5,10,11)^.

In addition to their canonical role in immune response, there is increasing evidence to support an extended role and functional diversification of AMPs ^(12)^. In particular, recent works are challenging the original view on AMPs as “killer molecules” by unravelling their participation in the “management” of commensal and mutualistic bacteria. For instance, the AMP production by epithelial cells was shown to regulate the beneficial gut microbiota composition and abundance in *Drosophila* ^(13)^ and the honey bee *Aphis mellifera* ^(14)^. Similarly, the compartmentalisation of immune related genes and fine-tuning of AMP expression in the different parts of the gut of the oriental fruit fly *Bactrocera dorsalis* results in the separation of diverse niches that favour the cultivation of beneficial microbes ^(15)^. Additionally, it was demonstrated in *Drosophila* that gene duplication and sequence convergence of the AMPs Diptericins could evolve diverse functions, resulting in a species-specific AMP repertoire needed for the adaptation to the occupied ecological niche and its associated microbial community ^(16,17)^.

Moreover, a function for AMPs in the “management” of beneficial symbionts has also been demonstrated in some cases of intracellular symbiosis from insects to plants ^(18,19)^. Hence, the renewed attention to beneficial animal-microbe interactions in the last decades and the constant refinement in understanding animal immunity ^(20,21)^ are converging towards an integrated view of AMPs that questions whether the term “antimicrobial” should evolve to take into consideration this new knowledge on their multiple functions, as proposed by Bosch *et al*. in this special issue ^(22)^.

Here, we will discuss the functions of AMPs in insects’ mutualistic association with bacteria, through a focus on the cereal weevil and its intracellular symbiont (endosymbiont) *Sodalis pierantonius*, with particular emphasis to AMPs’ functions in immune responses to pathogenic infection, control and maintenance of endosymbiosis, and during developmental processes. Cereal weevils are among the most advanced models in the field of immune adaptation to endosymbiotic constraints, and of particular significance to illustrate the duality of the immune response to both mutualistic symbionts and pathogens. Its binary association with a relatively recently acquired endosymbiont, the availability of host and endosymbiont genomic sequences and host genetic tools, as well as the possibility to obtain artificially non-symbiotic (aposymbiotic) weevils, provide appropriate conditions to tackle emerging questions in the field of immune interaction with microbiota. Here, we review the recent advances made on the role of AMPs in this model system and place them in perspective with discoveries in other models. We also present some novel experimental data that complete our understanding of the regulation of AMPs at metamorphosis.

## 1. The immune effector arsenal in a symbiotic insect encompasses a cocktail of AMPs

The innate immune response is the first line of defence against invasion by microbial pathogens ^(23)^. In insects, this response is divided into humoral defences that trigger the secretion of effector molecules by the fat-body and other tissues, and cellular defences, such as phagocytosis and encapsulation, that are mediated by haemocytes, the circulating innate immune cells ^(24)^.

The humoral defence has been broadly deciphered in the genetic model *Drosophila melanogaster* and was described to be regulated by two main signalling pathways, the Toll, and Immune deficiency (IMD) pathways, which regulate the expression of effector genes, including genes encoding AMPs, in response to infection ^(25,26)^. Both pathways are activated in response to the detection of peptidoglycan (PG), a cell wall component of bacteria. The Toll pathway is activated upon recognition by receptors Gram-Negative Binding Protein-1 (GNBP-1) and PeptidoGlycan Recognition Protein (PGRP)-SA of Lysine (Lys) type PG present in Gram-positive cocci bacteria. The IMD pathway is activated by receptors PGRP-LC and PGRP-LE upon recognition of diaminopimelic acid (DAP) type, present in Gram-negative bacteria and Gram-positive bacilli ^(24)^. However, extending the understanding of the innate immune mechanisms to other insects is important to define what are the conserved or acquired features of the immune system selected in these species through evolution and their adaptation to various environments, including the biotic environment. Importantly, an estimate of 15 to 20% of the insect species have evolved intracellular nutritional endosymbioses, in which the endosymbiont provides metabolites to the insect, hence allowing it to thrive on unbalanced diets, such as the mammalian blood, plant saps or cereal grains ^(27)^. Endosymbionts are transmitted vertically for thousands of host generations, leading to a long co-evolution of the host and bacterial partners. These associations are of great interest to address the question of how the once-infectious bacteria and the host immune system evolve in this context of mutualistic symbiosis.

Three species of *Sitophilus*, the granary weevil *S. granarius*, the rice weevil *S. oryzae* and the maize weevil *S. zeamais* are among the most important agricultural pests, causing extensive damage to cereal in fields and to stored grains. These coleopterans from the Curculionidae family are holometabolous insects undergoing complete metamorphosis. The life cycle of cereal weevils can be divided into four stages: egg, larva, pupa, and adult. The first three stages occur exclusively inside the cereal grain where the egg has been laid. Cereal weevils have evolved an obligate symbiotic relationship with the Gram-negative γ-proteobacterium *Sodalis pierantonius*, which complements their diet with amino acids, vitamins and co-factors ^(28–34)^. Endosymbionts are maintained within specific host cells, the bacteriocytes, which group into an organ, the bacteriome. One major evolutionary interest of the *Sitophilus/Sodalis* association relies on its relatively recent age, estimated to be established around 30,000 years ago ^(32,35–38)^. When compared to long-lasting endosymbionts, the genome of endosymbiont *S. pierantonius* is much less degenerated and reduced, with a size similar to the genomes of free-living enterobacteria ^(32,38)^. Interestingly, this genome encodes pathogenesis-related genes, including genes encoding Type Three Secretion System (TTSS), and genes necessary for cell wall synthesis, including DAP-type PG ^(32)^. Accordingly, *S. pierantonius* was shown to trigger a systemic expression of AMPs when experimentally injected inside *S. zeamais* ^(39)^, indicating it is able to be recognized by the host immune system when present extracellularly, possibly through the detection of its PG ^(40)^. Therefore, this association provides a useful framework to understand how endosymbiosis modulates the host immune system at a relatively “early” stage of the symbiotic partners’ co-evolution.

In order to complete our knowledge on immune pathways and effector-encoding genes in cereal weevils, we have recently sequenced *S. oryzae* genome and annotated the immune genes ^(34)^. On the basis of the mechanistic knowledge acquired in different species and in particular in *Drosophila*, this analysis showed that the Toll and IMD pathways are conserved in *S. oryzae* (**Figure 1**). For Toll pathway, only the genes encoding for the G protein-coupled receptor kinase 2 (Gprk2), Refractory to sigma P (Ref(2)P) and the transcription factor Dorsal-related immunity factor (Dif) appear to be lost, suggesting the pathway is solely relying on the transcription factor Dorsal for transcriptional activation of target genes. As for the IMD pathway, the genome encodes all the main and essential components of the pathway, with the notable exception of the intracellular receptor PGRP-LE, suggesting that the IMD pathway is functional but may lack the possibility to be activated by intracellular bacteria ^(34)^. Three main groups of immune effectors were identified in *S. oryzae*, namely AMPs, lysozymes and thaumatins ^(34)^. AMPs generally consist of twelve to fifty amino acids, with the hydrophobic part of the molecule generally covering more than 50% of amino acids residues. AMPs are divided into subgroups based on their amino acid composition and structure ^(7,41)^. Based on these features, sixteen sequences matching four different categories of AMPs were identified in *S. oryzae* by combining automated genome annotations and manual curations (**Figure 2; Supplementary table 1**). We found well-known peptides, including i) Defensin, characterised by six to eight conserved cysteine residues, a stabilising array of three or four disulphide bridges and three domains consisting in a flexible amino-terminal loop; ii) Cecropin, characterised by linear peptides with alpha-helix that lack cysteine residues; iii) Cathelicidin-like AMPs, a class of small, cationic peptides that play a crucial role in the innate immune system of many organisms, including humans and other vertebrates ^(42)^ and iv) Diptericins, Sarcotoxin, Coleoptericins and other members of the Glycine-rich AMPs, characterised by an over-representation of proline and/or glycine residues ^(34)^. Intriguingly, the genes *diptericin-3* and *diptericin-4*, located next to each other in the genome, encode identical peptides although they differ in their intronic regions, suggesting a recent duplication event. In light of the recent work on deciphering the evolution of *diptericins* in *Diptera* ^(16,43,44)^, it will be interesting to examine the Diptericin-encoding genes in closely related *Sitophilus* species, as well as a potential polymorphism in various populations of *S. oryzae*.

**Figure 1.**
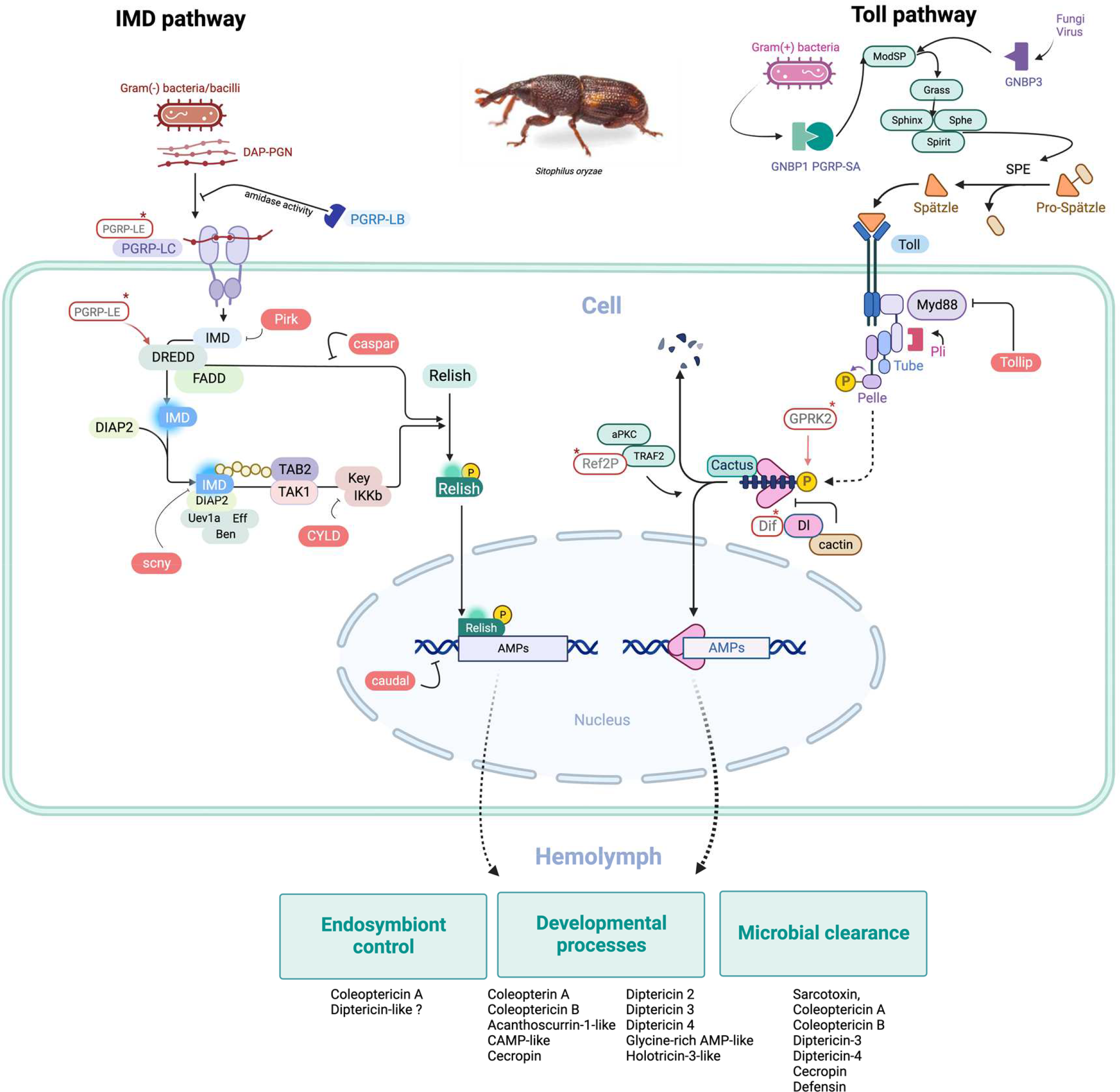
Schematic representation of IMD and Toll pathways in the cereal weevil *Sitophilus oryzae*. The immune signalling transduction pathways IMD and Toll activating AMP-encoding genes in *S. oryzae* were generated with Biorender (https://biorender.com/). All members of these pathways were manually verified by blast (Parisot *et al*., 2021). The name of each member corresponds to *Drosophila* protein names. Missing members are shown in empty rectangles in red. IMD and Toll pathways trigger the expression of AMP that can participate to different functions in *S. oryzae*, including endosymbiont control, developmental processes, and microbial clearance.

**Figure 2.**
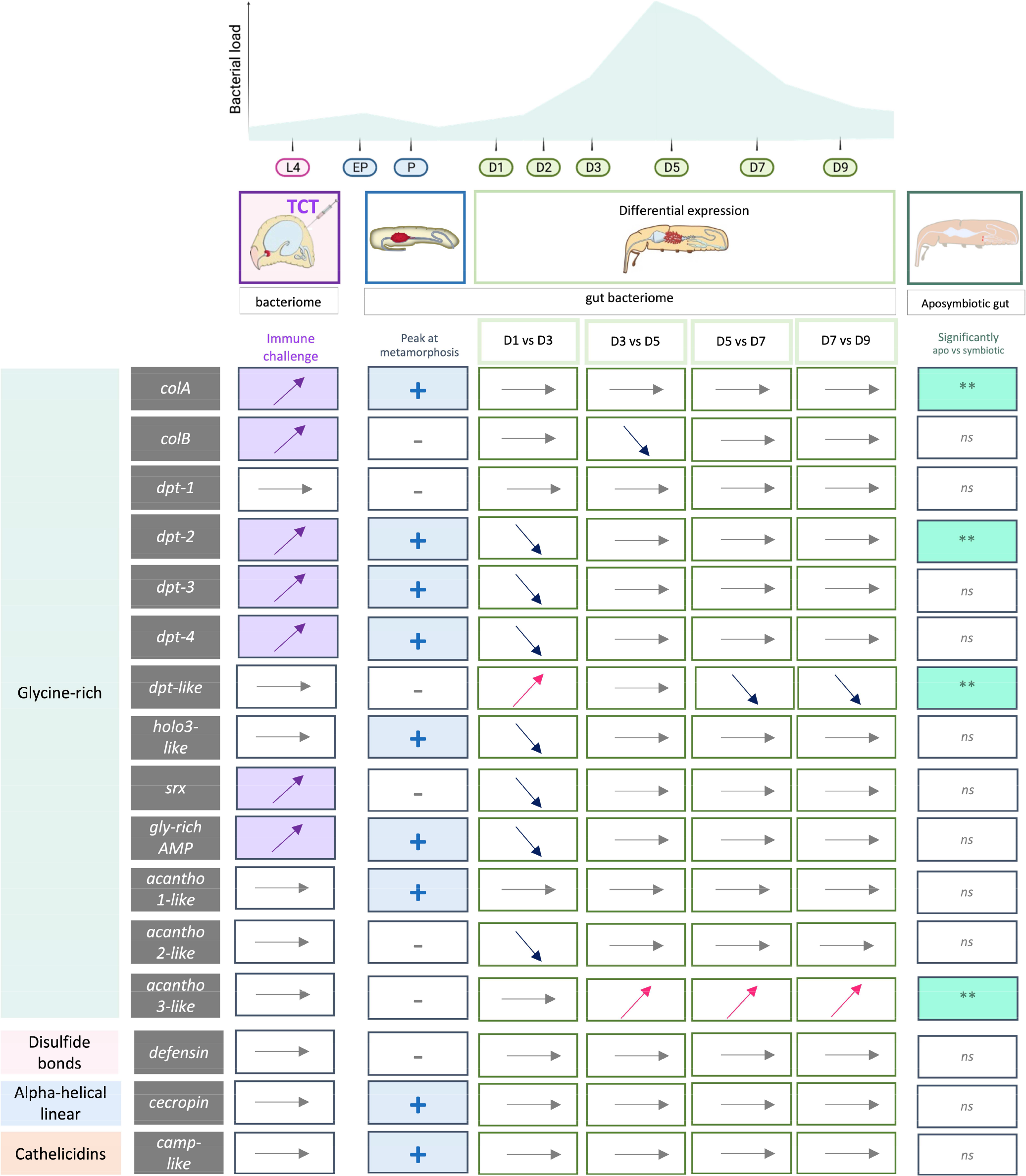
Differential expression profile of AMPs in symbiotic and aposymbiotic weevils during development or under immune challenge. Genes were considered as differentially expressed when the adjusted P-value was lower than 0.05 and the absolute Log2 of fold change was greater than 0.5 in fourth-instar larvae (L4), early pupal stage (EP), pupal stage (P), late pupal stage (LP) and day-one adult (D1) to day-nine adult (D9). In pupae (P), the sign plus in blue indicates a peak expression of AMP at metamorphosis. In the last column, the double asterisk indicates a significantly different expression of AMPs between symbiotic and aposymbiotic adult guts (from data published in Ferrarini *et al*., 2023).

## 2. AMPs in pathogenesis

The first historically identified function of AMPs is to fight bacterial intruders. In the last decade, studies performed in *S. oryzae* and *S. zeamais* have deciphered the immune response of these cereal weevil species against infection and identified the AMP effectors involved. These weevils demonstrate a potent immune response when infected: the expression of *coleoptericin A (colA), coleoptericin B (colB), sarcotoxin, diptericin-3, diptericin-4, cecropin* and *defensin* was shown to be systemically induced in *S. oryzae* larvae, with a peak of expression at 12 h following injection of either Gram-positive (*Micrococcus luteus*) or Gram-negative (*Dickeya dadantii*) bacteria ^(45)^. We further investigated how weevils respond to Gram-negative bacteria infection for better understanding weevil’s immune adaptation to the Gram-negative endosymbiont *S. pierantonius* ^(40)^. Maire *et al*. showed that both *S. oryzae* and *S. zeamais*’s immune system could be activated by purified *Escherichia coli* tracheal cytotoxin (TCT), which is a monomer of DAP-type PG that was identified as a minimal and very efficient elicitor of the IMD pathway in *D. melanogaster* ^(40,46)^. Injection of TCT has the advantage of avoiding any potential interference of living bacteria with host immunity. TCT injection systemically induced the expression of *colA, colB* and *sarcotoxin* in *S. zeamais* ^(40)^. RNA interference (RNAi) experiments further confirmed that this systemic expression of weevil’s AMPs is Imd and Relish dependent in both *S. oryzae* and *S. zeamais* ^(40)^, similarly to what was demonstrated in *Drosophila* ^(24)^ and other holometabolous insects, including the yellow fever mosquito *Aedes aegypti* ^(47)^, *A. mellifera* ^(48)^, pollen beetle *Meligethes aeneus* ^(49)^, greater wax moth *Galleria mellonella* ^(50)^, red flour beetle *Tribolium castaneum* ^(51)^ and mealworm *Tenebrio molitor* ^(52)^.

Under standard symbiotic conditions (*i*.*e*. absence of infection by free-living bacteria), the bacteriome expresses a specific immune program. Despite the massive presence of endosymbionts, this organ was shown to not express strongly most of the genes encoding AMPs tested but one, named *coleoptericin A* (*colA*) in *S. zeamais* ^(39)^. Nevertheless, the bacteriome was found to be immune responsive when both *S. oryzae* and *S. zeamais* were challenged with infectious bacteria. AMP-encoding genes such as *colA, colB, sarcotoxin, diptericin-3, diptericin-4, cecropin* and *defensin* were shown to be locally expressed in the bacteriome after an immune challenge, although at a lower amount compared to the systemic response ^(40,45)^. Moreover, dual RNAseq on the bacteriome of *S. oryzae* after TCT injection further confirmed that the bacteriome actively participates in the immune response by the upregulation of 11 genes, among which 7 were AMP-encoding genes, namely c*olA, colB, sarcotoxin, gly-rich AMP-like* and three *diptericins* (**Figure 2**) ^(53)^. Strikingly, the endosymbiont load was not affected neither by pathogenic infection itself nor by the systemic immune response triggered by the infection ^(45)^. The dual RNAseq revealed that the endosymbionts did not undergo any transcriptional changes in response to the TCT injection, suggesting that the potential threat from AMPs went unnoticed by the endosymbiont ^(53)^. Immunohistochemical observations reported in Ferrarini *et al*. showed that even though the bacteriome produces AMPs upon an immune challenge, their final location is outside the bacteriome, hence providing physical separation from endosymbionts ^(53)^.

The physical separation of endosymbionts by compartmentalisation in the bacteriome not only limits their exposure to the host humoral and cellular immune responses but also allows the expression of a tissue-specific immune program ^(19,39)^. Such an “innovative” differentiation of specific cells and organ permits the execution of a specific immune program by synthesising immune effectors that can either be secreted to fight against pathogens or target endosymbionts within the bacteriocytes ^(54)^.

## 3. Immune evolution in the context of endosymbiosis

Maintenance of obligatory nutritional endosymbiosis represents a “cohabitation challenge” for the host and bacteria. Over co-evolution, in ancient endosymbioses, a degeneration of the bacterial genome is observed that leads to the loss of genes that are not essential for the intracellular life style of the bacteria ^(55,56)^. Genes encoding proteins involved in pathogenesis, including cell-surface components (lipopolysaccharides, phospholipids), flagellum, TTSS, are often pseudogenised or lost ^(38,57,58)^, which decrease the infectivity and immunogenicity of endosymbionts. As mentioned above, this is not the case in *S. pierantonius*, which has more recently replaced a previous endosymbiont in the association with *Sitophilus* spp. ^(32,36–38)^. *S. pierantonius* genome retains similar genomic features to free-living bacteria ^(32,58)^. Hence S*itophilus/Sodalis* association offers the opportunity to address whether the host genome and especially the genes encoding immune system components are also shaped by the host-endosymbiont co-evolution. For example, we have shown that the *pgrp-lb* gene, known in other species to regulate the immune response to infection and gut commensal ^(59,60)^, encodes isoforms in *S. oryzae* and *S. zeamais* that ensure host immune homeostasis in the context of endosymbiosis ^(61)^.

Another example of host genome adaptation with regard to endosymbiosis is the AMP ColA ^(19)^. In more details, *colA* gene is particularly highly expressed in the bacteriome of *S. oryzae* and *S. zeamais* ^(39)^ and RNAi against *colA* demonstrated that ColA prevents endosymbiont escape from the bacteriocytes ^(19)^. Moreover, immunogold staining combined with electron microscopy observations showed that ColA targets the endosymbiont cytosol, and far-Western blotting unravelled that ColA interacts with the bacterial chaperonin GroEL ^(19)^. Remarkably, ColA was shown to inhibit the bacterial cell division without interfering with DNA replication, resulting in the formation of polyploid, metabolically-active, gigantic bacterial cells ^(19)^. In addition to ColA, another coleoptericin, called Coleoptericin B (ColB), was identified in *S. oryzae*’s genome. Despite having 42% amino acid sequence identity with ColA, and similar bactericidal activity against the Gram-positive (*Micrococcus luteus*) and Gram-negative (*Escherichia coli*) free-living bacteria, ColB does not interact with *S*.*pierantonius*’s GroEL ^(19)^. The gene encoding ColB is not expressed in the bacteriome under standard symbiotic conditions, but only after external immune challenges mimicking systemic infection with free-living bacteria ^(19)^. This suggests distinct functions for these two AMPs in weevil immunity ^(19)^. Hence, among the Coleoptericin family, the AMP ColA seems to have acquired a specific function in weevil-endosymbiont association, where this AMP interferes with bacterial cytokinesis leading to bacterial cell filamentation, and prevent endosymbionts ‘escape from bacteriocytes. Interestingly, ColA has also retained a canonical AMP function in insect immunity, demonstrated by its bactericidal activity and the systemic upregulation of its expression during infection. The progress made in RNAseq approaches have recently opened up the possibilities to analyse gene expression along the life cycle and in relation with symbiosis much more easily and at the whole transcriptomic scale. Analysing dual RNAseq data acquired along 12 stages of *S. oryzae* and comparing with RNAseq performed in aposymbiotic adults suggest a variety of functions for cereal weevil’s AMP, including a function in the control of endosymbiosis that would not be solely restricted to ColA (**Figure 2**) ^(62)^. Indeed, the expression profile of several AMP genes seems to parallel the dynamic changes in bacterial load during development. For example, *cathelicidin-like antimicrobial protein, acanthoscurrin-1-like* and *diptericin-like* AMPs are, as *colA*, highly expressed in the bacteriome at larval stages ^(62)^. Moreover, the expression level of gene encoding a novel putative intracellular AMP, the Diptericin-like AMP, was found to be highly correlated with bacterial load during weevil development ^(62)^. Functional analyses are required to assess potential symbiosis-related functions of these AMPs. Other AMPs were shown to be abundantly expressed in bacteriocytes in other insect species, including the Bacteriocyte-specific Cysteine-Rich (BCR) peptides *A. pisum* ^(63,64)^, but their function with regards to symbiosis remains to be investigated.

It is fascinating to see that AMPs were also shown to target bacterial cytokinesis while preserving DNA replication in plant-bacterial nutritional symbiotic associations ^(18,65)^. In the symbiosis of legume plants with the nitrogen-fixing endosymbiotic rhizobia, a large number of AMPs, called Nodule-specific Cysteine-Rich peptides, modify rhizobia into elongated bacteria that are highly efficient in nitrogen fixation ^(18)^. This process irreversibly renders the bacteria unable to resume growth or reproduce by interrupting the regular cycle of replication and division, resulting in polyploid cells ^(18,65)^, similarly to *S. pierantonius* in *Sitophilus* spp. under the action of ColA. Likewise, in *Alnus-Frankia* endosymbiosis, nodules produce Defensin-like AMPs, among which Ag5 that was demonstrated to target *Frankia* cells and increases membrane permeability, suggesting a function in metabolite exchange between endosymbionts and host cells ^(66,67)^. Thus, AMPs could favour the production of metabolites, through polyploidy of bacteria, as well as metabolite exchanges, through increased permeability of the endosymbiont membrane ^(67)^. This points to a diversification of the AMPs’ mode of action, from the clearance of pathogens to the facilitation of nutrient exchanges in nutritional endosymbiosis. In this regard, several other mechanisms through which AMPs target pathogens could be reused in the context of symbiosis. AMPs can directly act on intracellular target molecules, including DNA, RNA and proteins, they can inhibit cell wall synthesis, nucleic acid and protein synthesis, or enzymatic activity ^(4)^. Moreover, AMPs can inhibit bacterial cytokinesis, interfere with division machinery or regulators of the bacterial cell cycle ^(68,69)^. These examples are suggesting an arsenal of modes of action that can be exploited and selected during long-term host-endosymbiont co-evolution.

## 4. AMPs’ involvement during developmental processes

An increasing number of evidences indicate that AMPs play a role during developmental processes, either during embryonic or post-embryonic development (**Table 1** and references within). Post-embryonic development is particularly important in insects such as cereal weevils experiencing complete metamorphosis (holometabolism), for which the transition between the larval and adult stages is achieved through the nonfeeding pupal stage, in which tissues are *de novo* synthesised and larval tissues are replaced, degenerated or undergo remodelling ^(70)^. This massive reorganisation of the organism is synchronised and coordinated by the steroid hormone ecdysone ^(71)^.

**Table 1.** AMPs peak at metamorphosis in different holometabolous insect lineages.

| Order, Development | Insect | AMP | Developmental stage | Anticipated function | References |
| --- | --- | --- | --- | --- | --- |
| Diptera | Flesh fly<br>(*Sarcophaga peregrina*) | Defensin, Sarcotoxin | Embryo, Pupae | Growth factor, ontogenesis | (73,74) |
| Lepidoptera | Tobacco hornworm<br>(*Manduca sexta*) | Attacins, Cecropins,<br>Gloverins, Lebocins, Dipausin | Pupae | Pathogen clearance/control<br>during moulting | (75,76) |
| Lepidoptera | Diamondback moth<br>(*Plutella xylostella*) | Gloverins | Pupae | Pathogen clearance/control<br>during moulting | (77) |
| Lepidoptera | Silkworm pupae<br>(*Bombyx mori*) | Attacin, Cecropins, Gloverins,<br>Defensin, Hemolin, Lebocin,<br>Moricin | Pupae | Pathogen clearance/control<br>during moulting | (78,79) |
| Coleoptera | Flour beetle pupae<br>(*Tribolium castaneum*) | Cecropin,<br>Defensin | Pupae | Pathogen clearance/control<br>during moulting | (81) |
| Diptera | *Drosophila melanogaster* | Drosomycin, Drosomycin-like | Pupae | Pathogen clearance/control<br>during moulting | (84) |

In *S. oryzae*, the larval bacteriome dissociates during complete metamorphosis of the insect, and bacteriocytes migrate before forming discrete repeated clusters of cells along the midgut, from where endosymbionts *de novo* infect nearby precursor cells (**Figure 3A**) ^(72)^. It was speculated that this infection could orientate the differentiation of these precursor cells into new bacteriocytes ^(72)^. On the endosymbiont side, flagellum (operons *fli* and *flh*) and TTSS-encoding genes (operons *ssa* and *sseE*) were found transcriptionally upregulated during metamorphosis, suggesting their involvement in the *de novo* infection process ^(58,62,72)^. Concomitantly, in *S. oryzae*, ten AMP-encoding genes were found to be transcriptionally upregulated when performing RNAseq on the dissected gut and bacteriomes at pupal stage, namely *colA, colB, acanthoscurrin-1-like, cathelicidin-like antimicrobial protein, cecropin, diptericin 2, diptericin 3, diptericin 4, glycine-rich AMP-like*, and *holotricin-3-like* (**Figure 2**) ^(62)^. At first sight, it is tempting to interpret this peak in AMP gene expression as the consequence of either the endosymbiont’s virulence gene expression, or their transient extracellular location while exiting the bacteriocytes. In this study, we aimed to clarify this point to address whether or not this peak of AMP expression is endosymbiont-dependent. Hence, we measured the transcriptomic steady-state level of *colA, colB, diptericin 2, diptericin 3, diptericin 4, glycine-rich AMP-like* and *sarcotoxin* in both symbiotic and aposymbiotic *S. oryzae* during metamorphosis. With the exception of *colA and sarcotoxin*, these novel data confirmed a significant induction of AMP expression at metamorphosis, but showed no significant differences between symbiotic and aposymbiotic weevils (**Figure 3B-G**). Thus, the peak in AMP gene expression during *S. oryzae* metamorphosis is independent of endosymbiont presence, and is therefore likely regulated by the insect’s developmental program and/or by metabolic changes occurring in the pupal non-feeding state.

**Figure 3.**
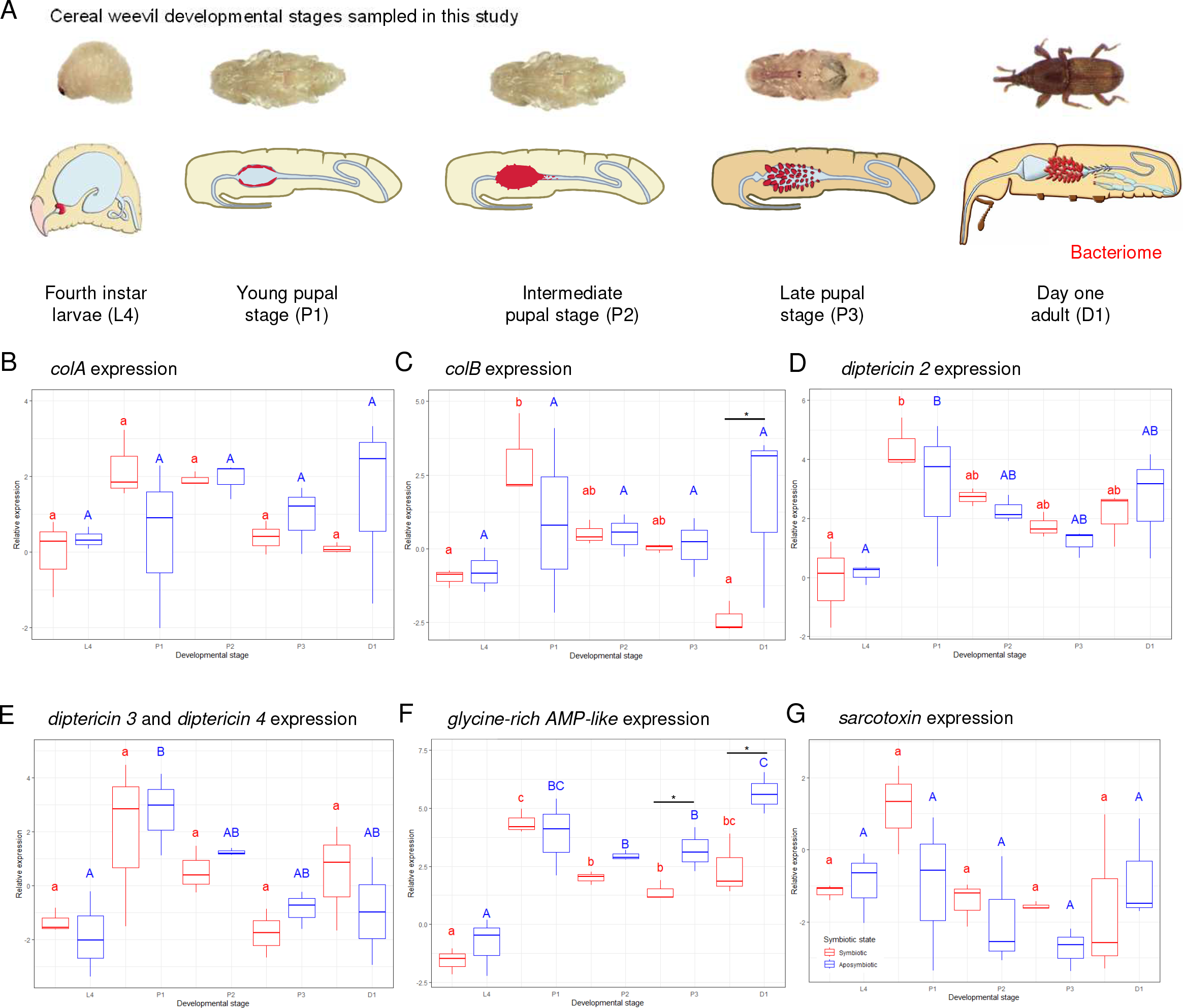
Expression of AMP-encoding genes during *Sitophilus oryzae* metamorphosis. Schematic representation of weevil developmental stages sampled in this experiment (A). Bacteriomes are illustrated in red. The Log^e^ transformation of relative expression of *colA* (B), *colB* (C), *diptericin 2* (D), *diptericin 3 and 4* (E), *glycine-rich AMP-like* (F) and *sarcotoxin* (G) in dissected guts of symbiotic (red) and aposymbiotic (blue) cereal weevils, measured at fourth-instar larvae (L4), young pupal stage (P1), intermediate pupal stage (P2), late pupal stage (P3) and day-one adult (D1). Mean and standard error values of three biological replicates, each consisting of five dissected guts, are presented for each developmental stage and symbiotic state. Different lowercase and uppercase letters indicate significant differences among developmental stages according to Tukey’s test (P ≤ 0.05). For each developmental stage star, symbols indicate significant differences in the pairwise comparisons between symbiotic and aposymbiotic insects according to Tukey’s test (P ≤ 0.05).

The first study reporting a peak in AMP expression at embryonic and pupal stages dates back to 1988, almost a decade after the first insect AMP description ^(9,73,74)^. Genes encoding Defensin and Sarcotoxin were found transcriptionally upregulated at the embryonic and pupal stages of the flesh fly (*Sarcophaga peregrina*), independently of experimental external stimuli ^(73,74)^. Since then, peaks of AMP expression at metamorphosis have been reported in many insects, either systemically or locally in the gut. For example, AMPs were found transcriptionally upregulated with other immune effectors in the pupae of several other lepidopteran species ^(75–80)^ and flour beetle *Tribolium castaneum* ^(81)^, in the absence of experimental infections (**Table 1**). These results, together with what we observed in cereal weevils, suggest that the expression of AMPs in the gut at metamorphosis is shared between a wild range of insects. Notably, when undergoing complete metamorphosis, insects face the challenge to replace the larval gut while concomitantly retaining beneficial symbiotic gut microbes in a controlled manner and avoiding infection by pathogens. Thus, the increased expression of AMPs at metamorphosis could be biologically relevant during the fragile period of moulting to prevent opportunistic microbes escaping from the gut lumen and potentially infecting insect tissues. Hence, AMP expression at metamorphosis could, at least in some species, be part of an ancient and conserved developmental program triggering “prophylactic” immunity independently of a current infection. To confirm this hypothesis, it would be interesting to test whether the AMP expression at metamorphosis is observed in axenic insects from the previous cited species.

Taken together, we can wonder how AMP gene expression could be regulated during metamorphosis, independently of bacterial presence ^(82,83)^. This question was recently genetically deciphered in the model *Drosophila*. Genes encoding Drosomycin (Drs), Drosomycin-like 2 (Drsl2) and Drosomycin-like 5 (Drsl5) are systemically activated during the metamorphosis of this insect, and show a peak of expression at pupal stage even in germ-free individuals^(84)^. In accordance with the bacteria-independent induction of the expression of these AMP-encoding genes, the NF-κB transcription factor Dif that is known to regulate *drs* under infection was not involved in this systemic expression of *drs* at metamorphosis. By driving the expression of a dominant negative form of the Ecdysone Receptor in the midgut or in the fat body, Nunes et al. ^(84)^, showed that *drsl2* local expression is regulated by ecdysone signalling in the gut at metamorphosis while *drs* systemic expression is regulated by ecdysone signalling in the fat-body, as *cecropin A1, defensin, metchnikowin* and *drsl3*. This effect of ecdysone on immune gene expression is in line with previous studies in *Drosophila* cell lines ^(82,83)^. Interestingly, Nunes *et al*. showed that *drs* was necessary to reduce bacteria remaining after imperfect larval gut purge at metamorphosis. Hence, ecdysone appears to not only coordinate the ontogenetic processes of metamorphosis but also to highest pre-emptive or “prophylactic” immune responses at metamorphosis in *Drosophila*, which had already been suggested for cellular responses ^(85)^. The double function of ecdysone as a master regulator of the metamorphosis and a positive modulator of the immune responses could be important to control gut microbiota composition and avoid opportunistic infections at this stage of morphological reconstruction ^(13,84,86,87)^.

In addition to ecdysone, other hormones have been proposed to participate in the regulation of AMPs at metamorphosis, including the neurohormone Bursicon, which regulates cuticle tanning in several insect species ^(88)^. In *A. aegypti* the relative expression of Bursicon subunits increased from very low in first instar larvae to high in pupae, before rapidly declining after adult emergence ^(89)^. In both *A. aegypti* and *D. melanogaster* the authors showed that overexpression of Bursicon subunits (α Burs and β Burs) induced AMP gene expression *via* the Relish transcription factor ^(89,90)^. These results suggest that Bursicon could function as a hormone transcriptionally regulating prophylactic immunity during times of heightened vulnerability, at metamorphosis and when newly formed adult cuticle is soft and easily wounded ^(89,90)^. *Bursicon* presents the same profile of high expression during metamorphosis, in *S. oryzae* ^(62)^. Further studies are required to confirm if AMP expression is ecdysone and/or Bursicon-dependent in cereal weevils.

To sum up, these examples provide evidence on the functional link between endocrine signalling and immunity ^(86)^, suggesting a role of developmental hormones in regulating the immune system over development and synchronising it with the specific immune needs of each stage.

## Conclusion

We aimed here to highlight some of the recent knowledge on AMPs’ functions in insects with a special focus on insect-microbe mutualistic symbiotic associations through the example of the *Sitophilus*-*Sodalis* association. We have shown that besides their role in immune responses, there is increasing evidence to support AMPs’ extended role and functional diversification, to the point that it could be pertinent to affiliate a new name for these small peptides. Throughout four very active decades of research on AMPs in insects, the diversification of the studied models and the availability of new genomic and transcriptomic techniques combined with targeted functional analyses continue unravelling the complexity of AMP functions in host-microbe co-evolution. Such studies provide guidelines for immunity research and disease management in organisms with adaptive immunity, including humans. In this regard, as outlined in several articles in this issue, further studies should decipher the spatiotemporal expression pattern, target range and specificity of AMPs ^(91–93)^. In particular, more attention should be dedicated to the synergistic and antagonistic effects among AMPs and how the expression of AMP-encoding genes is enhanced and controlled depending on the biotic and abiotic environment and developmental stage. The next step in the understanding of the complexity of AMP functions, regulations and evolution will likely come from the integration of population studies, as well as the integration of the multiple ecological factors that can modulate immunity and host-microbiome molecular interactions.

## Supporting information

Supplementary Table 2. List of primers used for RT-qPCR.

Supplementary Table 1. AMPs identified in the Sitophilus oryzae genome.

## Acknowledgments

We would like to acknowledge gratefully Hubert Charles for discussion about the statistics analysis, and the SymSIm team for assistance in preparing the samples and discussions. This work was funded by the ANR FOCuS (ANR-19-CE20-0010 -A. Zaidman-Rémy) and the Institut Universitaire de France (IUF -A. Zaidman-Rémy).

## Figure legends

**Supplementary Table 1. AMPs identified in the *Sitophilus oryzae* genome**.

**Supplementary Table 2. List of primers used for RT-qPCR**.

## References

1. Zasloff M. Antimicrobial peptides of multicellular organisms. Nature 2002; 415(6870): 389–95. 10.1038/415389a

2. Nes I. History, current knowledge, and future directions on bacteriocin research in lactic acid bacteria. Prokaryotic Antimicrobial Peptides Springer New York Dordrecht Heidelberg London; 2011. p. 3–12.

3. Brogden KA. Antimicrobial peptides: pore formers or metabolic inhibitors in bacteria? Nature Reviews Microbiology 2005; 3(3): 238–50. 10.1038/nrmicro1098

4. Li Y, Xiang Q, Zhang Q, Huang Y, Su Z. Overview on the recent study of antimicrobial peptides: Origins, functions, relative mechanisms and application. Peptides 2012; 37(2): 207–15. 10.1016/j.peptides.2012.07.001

5. Hanson MA, Dostálová A, Ceroni C, Poidevin M, Kondo S, Lemaitre B. Synergy and remarkable specificity of antimicrobial peptides in vivo using a systematic knockout approach. eLife 2019; 8: e44341. 10.7554/eLife.44341

6. Hanson MA, Kondo S, Lemaitre B. Drosophila immunity: the drosocin gene encodes two host defence peptides with pathogen-specific roles. Proceedings of the Royal Society B: Biological Sciences 2022; 289(1977): 20220773. 10.1098/rspb.2022.0773

7. Stączek S, Cytryńska M, Zdybicka-Barabas A. Unraveling the role of antimicrobial peptides in insects. International Journal of Molecular Sciences 2023; 24(6). 10.3390/ijms24065753

8. Zhang QY, Yan -B, Meng YM, Hong XY, Shao G, Ma JJ, et al. Antimicrobial peptides: mechanism of action, activity and clinical potential. Military Medical Research 2021; 8(1): 48. 10.1186/s40779-021-00343-2

9. Steiner H, Hultmark D, Engström Å, Bennich H, Boman HG. Sequence and specificity of two antibacterial proteins involved in insect immunity. Nature 1981; 292(5820): 246–8. 10.1038/292246a0

10. Zanchi C, Johnston PR, Rolff J. Evolution of defence cocktails: Antimicrobial peptide combinations reduce mortality and persistent infection. Molecular Ecology 2017; 26(19): 5334–43. 10.1111/mec.14267

11. Keshavarz M, Zanchi C, Rolff J. The effect of combined knockdowns of attacins on survival and bacterial load in Tenebrio molitor. Frontiers in Immunology 2023; 14. 10.3389/fimmu.2023.1140627

12. Hanson MA, Lemaitre B. New insights on Drosophila antimicrobial peptide function in host defense and beyond. Current Opinion in Immunology 2020; 62: 22–30. 10.1016/j.coi.2019.11.008

13. Marra A., Hanson M. A., Kondo S., Erkosar B., Lemaitre B. Drosophila antimicrobial peptides and lysozymes regulate gut microbiota composition and abundance. mBio 2021; 12(4): 10.1128/mbio.00824-21

14. Kwong WK, Mancenido AL, Moran NA. Immune system stimulation by the native gut microbiota of honey bees. Royal Society Open Science 2017; 4(2): 170003. 10.1098/rsos.170003

15. Yao Z, Cai Z, Ma Q, Bai S, Wang Y, Zhang P, et al. Compartmentalized PGRP expression along the dipteran Bactrocera dorsalis gut forms a zone of protection for symbiotic bacteria. Cell Reports 2022; 41(3). 10.1016/j.celrep.2022.111523

16. Hanson MA, Lemaitre B, Unckless RL. Dynamic evolution of antimicrobial peptides underscores trade-offs between immunity and ecological fitness. Frontiers in Immunology 2019; 10. 10.3389/fimmu.2019.02620

17. Hanson MA, Grollmus L, Lemaitre B. Ecology-relevant bacteria drive the evolution of host antimicrobial peptides in Drosophila. Science 2023; 381(6655): eadg5725. 10.1126/science.adg5725

18. Van de Velde W, Zehirov G, Szatmari A, Debreczeny M, Ishihara H, Kevei Z, et al. Plant peptides govern terminal differentiation of bacteria in symbiosis. Science 2010; 327(5969): 1122–6. 10.1126/science.1184057

19. Login FH, Balmand S, Vallier A, Vincent-Monégat C, Vigneron A, Weiss-Gayet M, et al. Antimicrobial peptides keep insect endosymbionts under control. Science 2011; 334(6054): 362–5. 10.1126/science.1209728

20. Eberl G. A new vision of immunity: homeostasis of the superorganism. Mucosal Immunology 2010; 3(5): 450–60. 10.1038/mi.2010.20

21. McFall-Ngai M, Heath-Heckman EAC, Gillette AA, Peyer SM, Harvie EA. The secret languages of coevolved symbioses: Insights from the Euprymna scolopes–Vibrio fischeri symbiosis. Seminars in Immunology 2012; 24(1): 3–8. 10.1016/j.smim.2011.11.006

22. Bosch T, McFall-Ngai M. A new lexicon in the age of microbiome research. Philosophical Transactions of the Royal Society B: Biological Sciences 2023;

23. Sheehan G, Garvey A, Croke M, Kavanagh K. Innate humoral immune defences in mammals and insects: The same, with differences? Virulence 2018; 9(1): 1625–39. 10.1080/21505594.2018.1526531

24. Lemaitre B, Hoffmann J. The host defense of Drosophila melanogaster. Annual Review of Immunology 2007; 25(1): 697–743. 10.1146/annurev.immunol.25.022106.141615

25. Cammarata-Mouchtouris A, Acker A, Goto A, Chen D, Matt N, Leclerc V. Dynamic regulation of NF-κB response in innate immunity: The case of the IMD pathway in Drosophila. Biomedicines 2022; 10(9). 10.3390/biomedicines10092304

26. Valanne S, Vesala L, Maasdorp MK, Salminen TS, Rämet M. The Drosophila Toll pathway in innate immunity: from the core pathway toward effector functions. The Journal of Immunology 2022; 209(10): 1817–25. 10.4049/jimmunol.2200476

27. Douglas AE. The microbial dimension in insect nutritional ecology. Functional Ecology 2009; 23(1): 38–47. 10.1111/j.1365-2435.2008.01442.x

28. Wicker C, Nardon P. Development responses of symbiotic and aposymbiotic weevils Sitophilus oryzae L. (Coleoptera, Curculionidae) to a diet supplemented with aromatic amino acids. Journal of Insect Physiology 1982; 28(12): 1021–4. 10.1016/0022-1910(82)90008-7

29. Wicker C. Differential vitamin and choline requirements of symbiotic and aposymbiotic S. oryzae (Coleoptera: Curculionidae). Comparative Biochemistry and Physiology Part A: Physiology 1983; 76(1): 177–82. 10.1016/0300-9629(83)90311-0

30. Gasnier-Fauchet F, Nardon P. Comparison of methionine metabolism in symbiotic and aposymbiotic larvae of Sitophilus oryzae L. (Coleoptera: Curculionidae) II. involvement of the symbiotic bacteria in the oxidation of methionine. Comparative Biochemistry and Physiology Part B 1986; 85(1): 251–4. 10.1016/0305-0491(86)90251-8

31. Gasnier-Fauchet F, Nardon P. Comparison of sarcosine and methionine sulfoxide levels in symbiotic and aposymbiotic larvae of two sibling species, Sitophilus oryzae L. and S. zeamais Mots. (Coleoptera: Curculionidae). Insect Biochemistry 1987; 17(1): 17–20. 10.1016/0020-1790(87)90138-7

32. Oakeson KF, Gil R, Clayton AL, Dunn DM, von Niederhausern AC, Hamil C, et al. Genome degeneration and adaptation in a nascent stage of symbiosis. Genome Biology and Evolution 2014; 6(1): 76–93. 10.1093/gbe/evt210

33. Vigneron A, Masson F, Vallier A, Balmand S, Rey M, Vincent-Monégat C, et al. Insects recycle endosymbionts when the benefit is over. Current Biology 2014; 24(19): 2267–73. 10.1016/j.cub.2014.07.065

34. Parisot N, Vargas-Chávez C, Goubert C, Baa-Puyoulet P, Balmand S, Beranger L, et al. The transposable element-rich genome of the cereal pest Sitophilus oryzae. BMC Biology 2021; 19(1): 241. 10.1186/s12915-021-01158-2

35. Heddi A, Charles H, Khatchadourian C, Bonnot G, Nardon P. Molecular characterization of the principal symbiotic bacteria of the weevil Sitophilus oryzae: A peculiar G + C content of an endocytobiotic DNA. Journal of Molecular Evolution 1998; 47(1): 52–61. 10.1007/PL00006362

36. Charles H, Heddi A, Rahbe Y. A putative insect intracellular endosymbiont stem clade, within the Enterobacteriaceae, inferred from phylogenetic analysis based on a heterogeneous model of DNA evolution. Comptes Rendus de l’Académie des Sciences - Series III - Sciences de la Vie 2001; 324(5): 489–94. 10.1016/S0764-4469(01)01328-2

37. Lefèvre C, Charles H, Vallier A, Delobel B, Farrell B, Heddi A. Endosymbiont phylogenesis in the Dryophthoridae weevils: Evidence for bacterial replacement. Molecular Biology and Evolution 2004; 21(6): 965–73. 10.1093/molbev/msh063

38. Clayton AL, Oakeson KF, Gutin M, Pontes A, Dunn DM, von Niederhausern AC, et al. A Novel human-infection-derived bacterium provides insights into the evolutionary origins of mutualistic insect–bacterial symbioses. PLOS Genetics 2012; 8(11): e1002990. 10.1371/journal.pgen.1002990

39. Anselme C, Pérez-Brocal V, Vallier A, Vincent-Monegat C, Charif D, Latorre A, et al. Identification of the weevil immune genes and their expression in the bacteriome tissue. BMC Biology 2008; 6(1): 43. 10.1186/1741-7007-6-43

40. Maire J, Vincent-Monégat C, Masson F, Zaidman-Rémy A, Heddi A. An IMD-like pathway mediates both endosymbiont control and host immunity in the cereal weevil Sitophilus spp. Microbiome 2018; 6(1): 6. 10.1186/s40168-017-0397-9

41. Wu Q, Patočka J, Kuča K. Insect Antimicrobial peptides, a mini review. Toxins 2018; 10(11). 10.3390/toxins10110461

42. Blower RJ, Popov SG, van Hoek ML. Cathelicidin peptide rescues G. mellonella infected with B. anthracis. Virulence 2018; 9(1): 287–93. 10.1080/21505594.2017.1293227

43. Hanson MA, Hamilton PT, Perlman SJ. Immune genes and divergent antimicrobial peptides in flies of the subgenus Drosophila. BMC Evolutionary Biology 2016; 16(1): 228 10.1186/s12862-016-0805-y

44. Unckless RL, Howick VM, Lazzaro BP. Convergent balancing selection on an antimicrobial peptide in Drosophila. Current Biology 2016; 26(2): 257–62. 10.1016/j.cub.2015.11.063

45. Masson F, Vallier A, Vigneron A, Balmand S, Vincent-Monégat C, Zaidman-Rémy A, et al. Systemic infection generates a local-like immune response of the bacteriome organ in insect symbiosis. Journal of Innate Immunity 2015; 7(3): 290–301. 10.1159/000368928

46. Stenbak CR, Ryu J-H, Leulier F, Pili-Floury S, Parquet C, Hervé M, et al. Peptidoglycan molecular requirements allowing detection by the Drosophila immune deficiency pathway. The Journal of Immunology 2004; 173(12): 7339–48. 10.4049/jimmunol.173.12.7339

47. Shin SW, Kokoza V, Lobkov I, Raikhel AS. Relish-mediated immune deficiency in the transgenic mosquito Aedes aegypti. Proceedings of the National Academy of Sciences of the United States of America 2003; 100(5): 2616–21.

48. Schlüns H, Crozier RH. Relish regulates expression of antimicrobial peptide genes in the honeybee, Apis mellifera, shown by RNA interference. Insect Molecular Biology 2007; 16(6): 753–9. 10.1111/j.1365-2583.2007.00768.x

49. Vogel H, Badapanda C, Knorr E, Vilcinskas A. RNA-sequencing analysis reveals abundant developmental stage-specific and immunity-related genes in the pollen beetle Meligethes aeneus. Insect Molecular Biology 2014; 23(1): 98–112. 10.1111/imb.12067

50. Sarvari M, Mikani A, Mehrabadi M. The innate immune gene Relish and Caudal jointly contribute to the gut immune homeostasis by regulating antimicrobial peptides in Galleria mellonella. Developmental & Comparative Immunology 2020; 110: 103732. 10.1016/j.dci.2020.103732

51. Choi B, Park W-R, Kim Y-J, Mun S, Park S-J, Jeong J-H, et al. Nuclear receptor estrogen-related receptor modulates antimicrobial peptide expression for host innate immunity in Tribolium castaneum. Insect Biochemistry and Molecular Biology 2022; 148: 103816. 10.1016/j.ibmb.2022.103816

52. Ko HJ, Patnaik BB, Park KB, Kim CE, Baliarsingh S, Jang HA, et al. TmIKKε is required to confer protection against Gram-negative bacteria, E. coli by the regulation of antimicrobial peptide production in the Tenebrio molitor fat body. Frontiers in Physiology 2022; 12. 10.3389/fphys.2021.758862

53. Ferrarini MG, Dell’Aglio E, Vallier A, Balmand S, Vincent-Monégat C, Hughes S, et al. Antimicrobial peptide secretion protects endosymbionts from bacteriome autoimmunity in insects. Immunology; 10.1101/2022.03.18.484386

54. Heddi A, Zaidman-Rémy A. Endosymbiosis as a source of immune innovation. Comptes Rendus Biologies 2018; 341(5): 290–6. 10.1016/j.crvi.2018.03.005

55. Moya A, Peretó J, Gil R, Latorre A. Learning how to live together: genomic insights into prokaryote–animal symbioses. Nature Reviews Genetics 2008; 9(3): 218–29 10.1038/nrg2319

56. Chong RA, Moran NA. Evolutionary loss and replacement of Buchnera, the obligate endosymbiont of aphids. Nature 2018; 12(3): 898–908 10.1038/s41396-017-0024-6

57. Shigenobu S, Watanabe H, Hattori M, Sakaki Y, Ishikawa H. Genome sequence of the endocellular bacterial symbiont of aphids Buchnera sp. APS. Nature 2000; 407(6800): 81–6 10.1038/35024074

58. Dale C, Plague GR, Wang B, Ochman H, Moran NA. Type III secretion systems and the evolution of mutualistic endosymbiosis. Proceedings of the National Academy of Sciences of the United States of America 2002; 99(19): 12397–402.

59. Zaidman-Rémy A, Hervé M, Poidevin M, Pili-Floury S, Kim M-S, Blanot D, et al. The Drosophila Amidase PGRP-LB Modulates the Immune Response to Bacterial Infection. Immunity 2006; 24(4): 463–73. 10.1016/j.immuni.2006.02.012

60. Charroux B, Capo F, Kurz CL, Peslier S, Chaduli D, Viallat-lieutaud A, et al. Cytosolic and secreted peptidoglycan-degrading enzymes in Drosophila respectively control local and systemic immune responses to microbiota. Cell Host & Microbe 2018; 23(2): 215–228.e4. 10.1016/j.chom.2017.12.007

61. Maire J, Vincent-Monégat C, Balmand S, Vallier A, Hervé M, Masson F, et al. Weevil pgrp-lb prevents endosymbiont TCT dissemination and chronic host systemic immune activation. Proceedings of the National Academy of Sciences 2019; 116(12): 5623–32. 10.1073/pnas.1821806116

62. Ferrarini, Mariana Galvão, Vallier, Agnès, Carole Vincent-Monégat, Elisa Dell’Aglio, Benjamin Gillet, Sandrine Hughes, et al. Coordination of host and endosymbiont gene expression governs endosymbiont growth and elimination in the cereal weevil Sitophilus spp. bioRxiv 2023; 10.1101/2023.04.03.535335

63. Shigenobu S, Stern DL. Aphids evolved novel secreted proteins for symbiosis with bacterial endosymbiont. Proceedings of the Royal Society B: Biological Sciences 2013; 280(1750): 20121952. 10.1098/rspb.2012.1952

64. Loth K, Parisot N, Paquet F, Terrasson H, Sivignon C, Rahioui I, et al. Aphid BCR4 structure and activity uncover a new defensin peptide superfamily. International Journal of Molecular Sciences 2022; 23(20). 10.3390/ijms232012480

65. Mergaert P, Uchiumi T, Alunni B, Evanno G, Cheron A, Catrice O, et al. Eukaryotic control on bacterial cell cycle and differentiation in the Rhizobium– legume symbiosis. Proceedings of the National Academy of Sciences of the United States of America 2006; 103(13): 5230–5. 10.1073/pnas.0600912103

66. Carro L, Pujic P, Alloisio N, Fournier P, Boubakri H, Hay AE, et al. Alnus peptides modify membrane porosity and induce the release of nitrogen-rich metabolites from nitrogen-fixing Frankia. Nature 2015; 9(8): 1723–33. 10.1038/ismej.2014.257

67. Mergaert P, Kikuchi Y, Shigenobu S, Nowack ECM. Metabolic integration of bacterial endosymbionts through antimicrobial peptides. Trends in Microbiology 2017; 25(9): 703–12. 10.1016/j.tim.2017.04.007

68. Tiricz Hilda, Szűcs Attila, Farkas Attila, Pap Bernadett, Lima Rui M., Maróti Gergely, et al. Antimicrobial nodule-specific cysteine-rich peptides induce membrane depolarization-associated changes in the transcriptome of Sinorhizobium meliloti. Applied and Environmental Microbiology 2013; 79(21): 6737–46. 10.1128/AEM.01791-13

69. Mergaert P. Role of antimicrobial peptides in controlling symbiotic bacterial populations. Natural Product Reports 2018; 35(4): 336–56. 10.1039/C7NP00056A

70. Rolff J, Johnston PR, Reynolds S. Complete metamorphosis of insects. Philosophical Transactions of the Royal Society B: Biological Sciences 2019; 374(1783): 20190063. 10.1098/rstb.2019.0063

71. Thummel CS. Molecular mechanisms of developmental timing in C. elegans and Drosophila. Developmental Cell 2001; 1(4): 453–65. 10.1016/S1534-5807(01)00060-0

72. Maire J, Parisot N, Galvao Ferrarini M, Vallier A, Gillet B, Hughes S, et al. Spatial and morphological reorganization of endosymbiosis during metamorphosis accommodates adult metabolic requirements in a weevil. Proceedings of the National Academy of Sciences of the United States of America 2020; 117(32): 19347–58. 10.1073/pnas.2007151117

73. Matsuyama K, Natori S. Molecular cloning of cDNA for sapecin and unique expression of the sapecin gene during the development of Sarcophaga peregrina. Journal of Biological Chemistry 1988; 263(32): 17117–21 10.1016/S0021-9258(18)37506-9

74. Nanbu R, Nakajima Y, Ando K, Natori S. Novel peature of expression of the sarcotoxin IA gene in development of Sarcophaga peregrina. Biochemical and Biophysical Research Communications 1988; 150(2): 540–4 10.1016/0006-291X(88)90427-5

75. Russell V, Dunn PE. Antibacterial proteins in the midgut of Manduca sexta during metamorphosis. Journal of Insect Physiology 1996; 42(1): 65–71. 10.1016/0022-1910(95)00083-6

76. He Y, Cao X, Li K, Hu Y, Chen Y, Blissard G, et al. A genome-wide analysis of antimicrobial effector genes and their transcription patterns in Manduca sexta. Insect Biochemistry and Molecular Biology 2015; 62: 23–37. 10.1016/j.ibmb.2015.01.015

77. Zhang Z, Zhu S, De Mandal S, Gao Y, Yu J, Zeng L, et al. Combined transcriptomic and proteomic analysis of developmental features in the immune system of Plutella xylostella during larva-to-adult metamorphosis. Genomics 2022; 114(4): 110381. 10.1016/j.ygeno.2022.110381

78. Wu S, Zhang X, He Y, Shuai J, Chen X, Ling E. Expression of antimicrobial peptide genes in Bombyx mori gut modulated by oral bacterial infection and development. Developmental & Comparative Immunology 2010; 34(11): 1191–8. 10.1016/j.dci.2010.06.013

79. Hou Y, Zhang Y, Gong J, Tian S, Li J, Dong Z, et al. Comparative proteomics analysis of silkworm hemolymph during the stages of metamorphosis via liquid chromatography and mass spectrometry. Proteomics 2016; 16(9): 1421–31. 10.1002/pmic.201500427

80. Vilcinskas A, Vogel H. Seasonal phenotype-specific transcriptional reprogramming during metamorphosis in the European map butterfly Araschnia levana. Ecology and Evolution 2016; 6(11): 3476–85 10.1002/ece3.2120

81. Critchlow JT, Norris A, Tate AT. The legacy of larval infection on immunological dynamics over metamorphosis. Philosophical Transactions of the Royal Society B: Biological Sciences 2019; 374(1783): 20190066. 10.1098/rstb.2019.0066

82. Flatt T, Heyland A, Rus F, Porpiglia E, Sherlock C, Yamamoto R, et al. Hormonal regulation of the humoral innate immune response in Drosophila melanogaster. Journal of Experimental Biology 2008; 211(16): 2712–24. 10.1242/jeb.014878

83. Rus F, Flatt T, Tong M, Aggarwal K, Okuda K, Kleino A, et al. Ecdysone triggered PGRP-LC expression controls Drosophila innate immunity. The EMBO Journal 2013; 32(11): 1626–38. 10.1038/emboj.2013.100

84. Nunes C, Koyama T, Sucena É. Co-option of immune effectors by the hormonal signalling system triggering metamorphosis in Drosophila melanogaster. PLOS Genetics 2021; 17(11): e1009916. 10.1371/journal.pgen.1009916

85. Regan JC, Brandão AS, Leitão AB, Mantas Dias ÂR, Sucena É, Jacinto A, et al. Steroid hormone signaling Is essential to regulate innate immune cells and fight bacterial infection in Drosophila. PLOS Pathogens 2013; 9(10): e1003720. 10.1371/journal.ppat.1003720

86. Nunes C, Sucena É, Koyama T. Endocrine regulation of immunity in insects. The FEBS Journal 2021; 288(13): 3928–47. 10.1111/febs.15581

87. Keith SA. Steroid hormone regulation of innate immunity in Drosophila melanogaster. PLOS Genetics 2023; 19(6): e1010782. 10.1371/journal.pgen.1010782

88. Fraenkel G, Hsiao C. Bursicon, a hormone which mediates tanning of the cuticle in the adult fly and other insects. Journal of Insect Physiology 1965; 11(5): 513–56. 10.1016/0022-1910(65)90137-X

89. Zhang H, Dong S, Chen X, Stanley D, Beerntsen B, Feng Q, et al. Relish2 mediates bursicon homodimer-induced prophylactic immunity in the mosquito Aedes aegypti. Scientific Reports 2017; 7(1): 43163. 10.1038/srep43163

90. An S, Dong S, Wang Q, Li S, Gilbert LI, Stanley D, et al. Insect neuropeptide bursicon homodimers induce innate immune and stress genes during molting by activating the NF-κB transcription factor relish. PLOS ONE 2012; 7(3): e34510. 10.1371/journal.pone.0034510

91. Klimovich, A., Bosch, T. Novel technologies uncover novel “anti”-microbial peptides in Hydra shaping the species-specific microbiome. Philosophical Transactions of the Royal Society B: Biological Sciences 2023;

92. Hanson MA. When the microbiome shapes the host: immune evolution implications for infectious disease. Philosophical Transactions of the Royal Society B: Biological Sciences 2023;

93. Hixon, B., Chen, R., Buchon, N. Innate immunity in Aedes mosquitoes: From pathogen resistance to shaping the microbiota. Philosophical Transactions of the Royal Society B: Biological Sciences 2023;

