## Supplementary Table 2. List of primers used for RT-qPCR. for "Cereal Weevil’s Antimicrobial Peptides: At the Crosstalk between Development, Endosymbiosis and Immune Response"

### Supplementary material

#### Material and methods

**Insect rearing and sampling.** *S. oryzae* weevils were reared on wheat grains in a climate chamber (27.5 °C, 70 % relative humidity, no light source). Aposymbiotic insects were obtained through heat-treatment [1,2]. *S. oryzae* were staged as follows: Forth-instar larvae (L4) are yellow, roundish, and possess a cephalic capsule. Young pupae (P1) are white; they lost the cephalic capsule; on their gut the mesenteric caeca have not yet developed. Intermediate pupae (P2) are white, with well-developed mesenteric caeca. Late pupae (P3) are brown and possess well-developed mesenteric caeca in the gut. Day-one adults (D1) are orange-coloured individuals unable to walk [3]. Guts from L4, P1, P2, P3 and D1 stage insects were dissected in diethylpyrocarbonate-treated Buffer A (25 mM KCl, 10 mM $MgCl_2$, 250 mM Sucrose, 35 mM Tris/HCl, pH 7.5). For RNA extraction five guts per developmental stage were pooled and stored at -80°C and each sampling was independently repeated three times.

**Total RNA extraction and reverse-transcription.** Total RNA extraction from dissected guts was performed with Ambion RNAqueous micro kit (AM1931) and genomic DNA was removed by DNase treatment, following the manufacturer's instructions. Total RNA concentration and quality were checked using a Nanodrop® spectrophotometer (Thermo Scientific) and agarose gel electrophoresis, respectively. Reverse transcription into the first strand cDNA was carried out using the iScript™ cDNA Synthesis Kit (Bio-Rad).

**Gene expression analysis by real-time quantitative reverse transcription RT-qPCR.** Transcript quantification was performed with a CFX Connect Real-Time detection system (Bio-Rad) using the LightCycler Fast Start DNA Master SYBR Green I kit (Roche Diagnostics). Data were calculated using the ratio of the target cDNA concentration to the geometric mean of two normalising gene concentrations: *cyclin-T1* (*CYCT1*; LOC115889772) and *protein phosphatase 1 regulatory subunit 12A* (*PP12A*; LOC115890036). Gene expression is therefore represented as the relative expression of the gene of interest compared with normalising genes. A complete list of primers used can be found in **Supplementary table 2**. PCR reactions were carried out as previously described [4].

**Statistical analysis.** Transcriptomic data were analysed with R software v4.3.1 (R Core Team, 2014). After validating log-transformed gene expression data for normal distribution (Shapiro-Wilk test, $P > 0.05$) and variance homogeneity of the data (Levene's tests, $P > 0.05$), significant differences in gene expression in the pairwise comparisons between symbiotic states and in multiple comparisons among developmental stages were assessed with the Tukey' test ($P \leq 0.05$).

### References


1. Nardon P. Obtention d'une souche aposymbiotique chez le charançon Sitophilus sasakii Tak : Différentes méthodes d'obtention et comparaison avec la souche symbiotique d'origine. Les Comptes Rendus de l'Académie des sciences 1973; 227D: 981-4.

2. Maire J, Parisot N, Galvao Ferrarini M, Vallier A, Gillet B, Hughes S, et al. Spatial and morphological reorganization of endosymbiosis during metamorphosis accommodates adult metabolic requirements in a weevil. Proceedings of the National Academy of Sciences 2020; 117(32): 19347-58. 10.1073/pnas.2007151117

3. Dell'Aglio Elisa, Lacotte Virginie, Peignier Sergio, Rahioui Isabelle, Benzaoui Fadéla, Vallier Agnès, et al. Weevil Carbohydrate Intake Triggers Endosymbiont Proliferation: A Trade-Off between Host Benefit and Endosymbiont Burden. mBio 2023; 14(2): e03333-22. 10.1128/mbio.03333-22

4. Maire J, Vincent-Monégat C, Masson F, Zaidman-Rémy A, Heddi A. An IMD-like pathway mediates both endosymbiont control and host immunity in the cereal weevil Sitophilus spp. Microbiome 2018; 6(1): 6. 10.1186/s40168-017-0397-9
