## Supplementary material for "Cereal Weevil’s Antimicrobial Peptides: At the Crosstalk between Development, Endosymbiosis and Immune Response"

| Accession number | Gene abbreviation | Gene whole name | Category of AMPs |
|---|---|---|---|
| LOC115874620 | *colA* | *coleoptericin A* | Glycine-rich AMPs |
| LOC115874703 | *colB* | *coleoptericin B* | Glycine-rich AMPs |
| LOC115877460 | *dpt-1* | *diptericin-1* | Glycine-rich AMPs |
| LOC115877462 | *dpt-2* | *diptericin-2* | Glycine-rich AMPs |
| LOC115877463 | *dpt-3* | *diptericin-3* | Glycine-rich AMPs |
| LOC115877465 | *dpt-4* | *diptericin-4* | Glycine-rich AMPs |
| LOC115877461 | *dpt-like partial* | *diptericin-like partial* | Glycine-rich AMPs |
| LOC115884869 | *holo3-like* | *holotricin-3-like* | Glycine-rich AMPs |
| LOC115888387 | *srx* | *sarcotoxin* | Glycine-rich AMPs |
| LOC115884866 | *gly-rich AMP* | *glycine-rich AMP like* | Glycine-rich AMPs |
| LOC115875079 | *acantho1-like* | *acanthoscurrin-1-like* | Glycine-rich AMPs |
| LOC115875523 | *acantho2-like* | *acanthoscurrin-2-like* | Glycine-rich AMPs |
| LOC115875524 | *acantho3-like* | *acanthoscurrin-3-like* | Glycine-rich AMPs |
| LOC115888712 | *defensin* | *defensin* | Disulfide bonds and beta-hairpin AMPs |
| LOC115891322 | *cecropin* | *cecropin* | Alpha-helical linear AMPs |
| LOC115886926 | *camp-like* | *cathelicidin-like antimicrobial protein* | Cathelicidins |
