## Supplementary Table 1. AMPs identified in the Sitophilus oryzae genome. for "Cereal Weevil’s Antimicrobial Peptides: At the Crosstalk between Development, Endosymbiosis and Immune Response"

| Accession number | Primer | Sequence (5'-3') |
|---|---|---|
| LOC115889772 | F *CYCT1* qPCR | CAGGACATGGGGCAAAGACTA |
| LOC115889772 | R *CYCT1* qPCR | TGGGAAGTCAGTGAAGGAGTG |
| LOC115890036 | F *PP12A* qPCR | TGCTCAAGGACGAGATTCGG |
| LOC115890036 | R *PP12A* qPCR | GCCAACCATTCTTTGGCGTC |
| LOC115874620 | F *colA* qPCR | GAATAGATACAACGGGGGTCA |
| LOC115874620 | R *colA* qPCR | CTACCATCTGACACTTCCTC |
| LOC115874703 | F *colB* qPCR | GCCCCCTAAGGCTCAAAAGA |
| LOC115874703 | R *colB* qPCR | ACTTGAGCTCTGGTGTTTCCA |
| LOC115877462 | F *diptericin 2* qPCR | GTGCTGTTCCATTGTTGGTG |
| LOC115877462 | R *diptericin 2* qPCR | ACCTCCACTATGGCTGTTGG |
| LOC115877463 | F *diptericin 3 and 4* qPCR | ACTCCGATTTCAAGCCGACA |
| LOC115877463 | R *diptericin 3 and 4* qPCR | ACTCCGATAACTCCTCCTCCT |
| LOC115877461 | F *diptericin-like AMP* qPCR | TACGGCAGCATTGGATTTCA |
| LOC115877461 | R *diptericin-like AMP* qPCR | TTTCCTTGCACCCTGGTTGA |
| LOC115884866 | F *glycine-rich AMP-like* qPCR | CAAAACCAAGATGAAATTTTTG |
| LOC115884866 | R *glycine-rich AMP-like* qPCR | CTGGGACTATCCCTGGAAGC |
| LOC115888387 | F *sarcotoxin* qPCR | AACCCTCCACTGTTGTAGAAG |
| LOC115888387 | R *sarcotoxin* qPCR | GGGCTTTGTATCTGCCAGTTG |
